# Independent Promoter Recognition by TcpP Precedes Cooperative Promoter Activation by TcpP and ToxR

**DOI:** 10.1101/2021.04.25.441279

**Authors:** A.L. Calkins, L.M. Demey, J.D. Karslake, E.D. Donarski, J.S. Biteen, V.J. DiRita

## Abstract

Cholera is a diarrheal disease caused by the Gram-negative bacterium *Vibrio cholerae*. To reach the surface of intestinal epithelial cells, proliferate, and cause disease, *V. cholerae* tightly regulates the production of virulence factors such as cholera toxin (*ctxAB*) and the toxin co-regulated pilus (*tcpA-F*). ToxT is directly responsible for regulating these major virulence factors while TcpP and ToxR indirectly regulate virulence factor production by stimulating *toxT* expression. TcpP and ToxR are membrane-localized transcription activators (MLTAs) required to activate *toxT* expression. To gain a deeper understanding of how MLTAs identify promoter DNA while in the membrane, we tracked the dynamics of single TcpP-PAmCherry molecules in live cells using photoactivated localization microscopy and identified heterogeneous diffusion patterns. Our results provide evidence that: 1) TcpP exists in three biophysical states (fast diffusion, intermediate diffusion, and slow diffusion); 2) TcpP transitions between these different diffusion states; 3) TcpP molecules in the slow diffusion state are interacting with the *toxT* promoter; and 4) ToxR is not essential for TcpP to localize the *toxT* promoter. These data refine the current model of cooperativity between TcpP and ToxR in stimulating *toxT* expression and demonstrate that TcpP locates the *toxT* promoter independent of ToxR.

## Introduction

The Gram-negative bacterium *Vibrio cholerae* infects millions of people each year, causing the diarrheal disease cholera resulting in ∼100,000 deaths annually ^1,2^, despite treatments available to combat infection, including vaccines, antibiotic therapy, and oral rehydration therapy ^3–10^. With changing climate and growing cases of antibiotic resistant *V. cholerae*, the number of annual cholera infections is projected to continue to increase ^11^. Thus, gaining deeper insight into the pathogenesis of *V. cholerae* will facilitate development of alternative methods of treatment, thereby reducing the global burden of cholera.

Upon ingestion, typically from contaminated water or food, *V. cholerae* colonizes the crypts of the villi in the distal portion of the small intestine and stimulates production of virulence factors essential for disease progression, such as the toxin co-regulated pilus and cholera toxin (TCP and CtxAB, respectively) ^12–17^. Transcription of *tcp* and *ctxAB* is directly activated by ToxT ^18–21^. Expression of *toxT* is highly regulated and positively stimulated by ToxR and TcpP, two membrane-localized transcription activators (MLTAs), which directly bind to the *toxT* promoter (*toxTpro*), with binding sites at −104 to −68 and −55 to −37, respectively ^18,22–28^. TcpP and ToxR are bitopic membrane proteins, each containing a cytoplasmic DNA-binding domain (within the PhoB and OmpR families respectively), a single transmembrane domain, and a periplasmic domain ^29^. ToxR appears to have an accessory role in *toxT* regulation. Evidence supporting the model that ToxR assists TcpP to *toxT* expression includes: i) TcpP binds downstream of ToxR, closer than ToxR to the putative RNA polymerase binding site on *toxTpro*; and ii) overexpression of TcpP results in ToxR-independent *toxT* transcription activation ^18,24,25,28^. Furthermore, we have previously measured the single-molecule dynamics of TcpP and noted that deletion of *toxR* decreases but does not eliminate the prevalence of TcpP-DNA binding events ^30^. As with other MLTAs, it remains unclear how TcpP and ToxR identify the *toxT* promoter while localized to the cytoplasmic membrane.

Signal transduction pathways in prokaryotes consist of one-component and two-component regulatory systems that manage cellular processes in response to extracellular information such as pH, temperature, chemical gradients, and nutrients ^31–33^. One-component regulatory systems combine their input and output functions in a single protein. MLTAs are a unique family of one-component regulators as they function from the cytoplasmic membrane, whereas the majority (∼97%) of one-component regulators are localized in the cytoplasm ^31^. These one-component membrane-localized regulators like TcpP and ToxR comprise a sensor domain and an output domain that are separated by a transmembrane domain. Some challenges emerge in understanding how MLTAs affect their function of activating transcription in response to external stimuli. For example, diffusion of these regulators is constrained to the cytoplasmic membrane. Additionally, the chromosome structure, which is not static, is known to influence association of a MLTA to its target sequence ^34–43^. How MLTAs locate their target sequences while bound to the membrane represents a major gap in our knowledge. Here, we investigated the subcellular single-molecule dynamics of TcpP-PAmCherry to understand how TcpP localizes to the *toxTpro* and to develop a general model for how MLTAs identify their DNA targets.

Our approach was to apply super-resolution single-molecule tracking (SMT) in living cells. Previous work demonstrated that TcpP molecules exhibit heterogeneous diffusion patterns ^30,44^. Here, we expand upon this earlier work to study the effect of specific mutations, that alter TcpP binding to DNA or the potential association of TcpP with ToxR, on TcpP subcellular mobility. By tracking the movement of TcpP-PAmCherry molecules within single living *V. cholerae* cells, we determined the distributions of the heterogeneous motions of TcpP, and detected changes in these diffusion coefficients in response to targeted genetic alterations. From this data, we identify three biophysical states (fast diffusion, intermediate diffusion, and slow diffusion), we propose a biological role of each state, and we suggest an alternative model of *toxT* activation where TcpP independently identifies the *toxTpro* prior to assistance from ToxR.

## Results

### Single-molecule tracking of TcpP-PAmCherry is useful to study promoter identification, but cannot probe regulated-intramembrane proteolysis

To investigate the dynamics of individual TcpP molecules, we generated a *V. cholerae* strain in which the wild type *tcpP* allele is replaced with one expressing TcpP fused at its C-terminus to a photoactivatable fluorescent protein, PAmCherry (*tcpP-PAmCherry*). Levels and activity of TcpP are controlled by a two-step proteolytic process known as regulated intramembrane proteolysis (RIP) ^45–47^. Under RIP-permissive conditions, the C-terminus of TcpP becomes sensitive to proteolysis by Tsp, a site-1 protease, and YaeL, a site-2 protease; this sensitivity results in the inability of the cell to activate *toxT* expression. Under non-RIP permissive conditions, TcpP is protected from RIP by TcpH ^45–47^.

We investigated whether we could assess RIP dynamics using single-molecule tracking. Like wild-type TcpP, TcpP-PAmCherry was sensitive to RIP in the absence of TcpH, indicated by lower levels of TcpP-PAmCherry in *tcpP-PAmCherryΔtcpH* relative to *tcpP-PAmCherry* (supplemental Figure 1). Secondly, in both *tcpP-PAmCherry* and *tcpP-PAmCherryΔtcpH* a smaller species of TcpP-PAmCherry was observed, referred to as TcpP-PAm* (supplemental Figure 1). A similar result has been observed for native TcpP in Δ*yaeL* cells and indicates RIP ^46^. Complementation of *tcpP-PAmCherryΔtcpH* with plasmid-encoded *tcpH* resulted in a band with the mass of native TcpP (∼29KDa), (supplemental Figure 2). These data indicate that TcpP-PAmCherry resists RIP in a TcpH-dependent fashion similar to native TcpP. As expected, native TcpP was not detected in the absence of TcpH. These data indicate that: 1) TcpP-PAmCherry is sensitive to RIP; 2) TcpH can protect TcpP-PAmCherry from RIP; and 3) addition of PAmCherry to the C-terminus of TcpP reduces RIP of TcpP-PAmCherry relative to TcpP. These conclusions are supported by similar levels of TcpA, CtxB, and *toxT* expression in *tcpP-PAmCherry* and *tcpP-PAmCherryΔtcpH* ^44^; (supplemental Figures 1 and 3). Notwithstanding the detectable levels of TcpP-PAmCherry on immunoblots of total proteins from *tcpP-PAmCherryΔtcpH*, we observed almost no TcpP-PAmCherry molecules in our single-molecule tracking experiments. As a result, we are unable to collect sufficient data to perform any analysis of *tcpP-PAmCherryΔtcpH* cells. Though we cannot determine how RIP influences TcpP-PAmCherry single-molecule dynamics, fusion of PAmCherry to the C-terminus of TcpP does not affect its ability to stimulate *toxT* expression (supplemental Figure 3). Therefore, TcpP-PAmCherry is an effective tool to understand how TcpP locates the *toxTpro* from its position in the membrane. Lastly, addition of PAmCherry to the C-terminus of TcpP does not affect the growth rate of *V. cholerae* (supplemental Figure 4).

### Baseline Dynamics of TcpP-PAmCherry

Single-Molecule Analysis by Unsupervised Gibbs sampling (SMAUG) characterizes the motion of molecules based on the collection of measured displacements (steps) in their single-molecule trajectories. SMAUG estimates the biophysical descriptors of a system by embedding a Gibbs sampler in a Markov Chain Monte Carlo framework. This non-parametric Bayesian analysis approach determines the most likely number of mobility states and the average diffusion coefficient of single molecules in each state, the population of each state, and the probability of transitioning between different mobility states over the course of a single trajectory ^44^. In our previous study, we determined that TcpP-PAmCherry molecules in *V. cholerae* cells transition between multiple biophysical states: fast diffusion, intermediate diffusion, and slow diffusion ^44^.

Here, we collected a new robust set of TcpP-PAmCherry tracking data in living *V. cholerae* cells (54,454 steps collected from 7601 trajectories) to further refine our analysis and to assign biochemical mechanisms to these biophysical observations (a sample of these tracks is shown in Figure 1b). Consistent with our previous results, we ascertained that TcpP-PAmCherry exists in three distinct states (slow diffusion, intermediate diffusion, and fast diffusion; blue, organge, and purple, respectively, in Figure 1c). Furthermore, we determined that TcpP-PAmCherry molecules do not freely transition between all the diffusion states: we observe that TcpP-PAmCherry molecules can transition between the fast state (purple) and the intermediate state (orange) and between the intermediate state (orange) and the slow state (blue) freely, but there is no significant probability of transitions directly from the fast diffusion state (purple) to the slow diffusion state (blue) on successive steps (Figure 1d). Thus, the intermediate diffusion state represents a critical biochemical intermediate between the slow and fast diffusion states.

**Figure 1:**
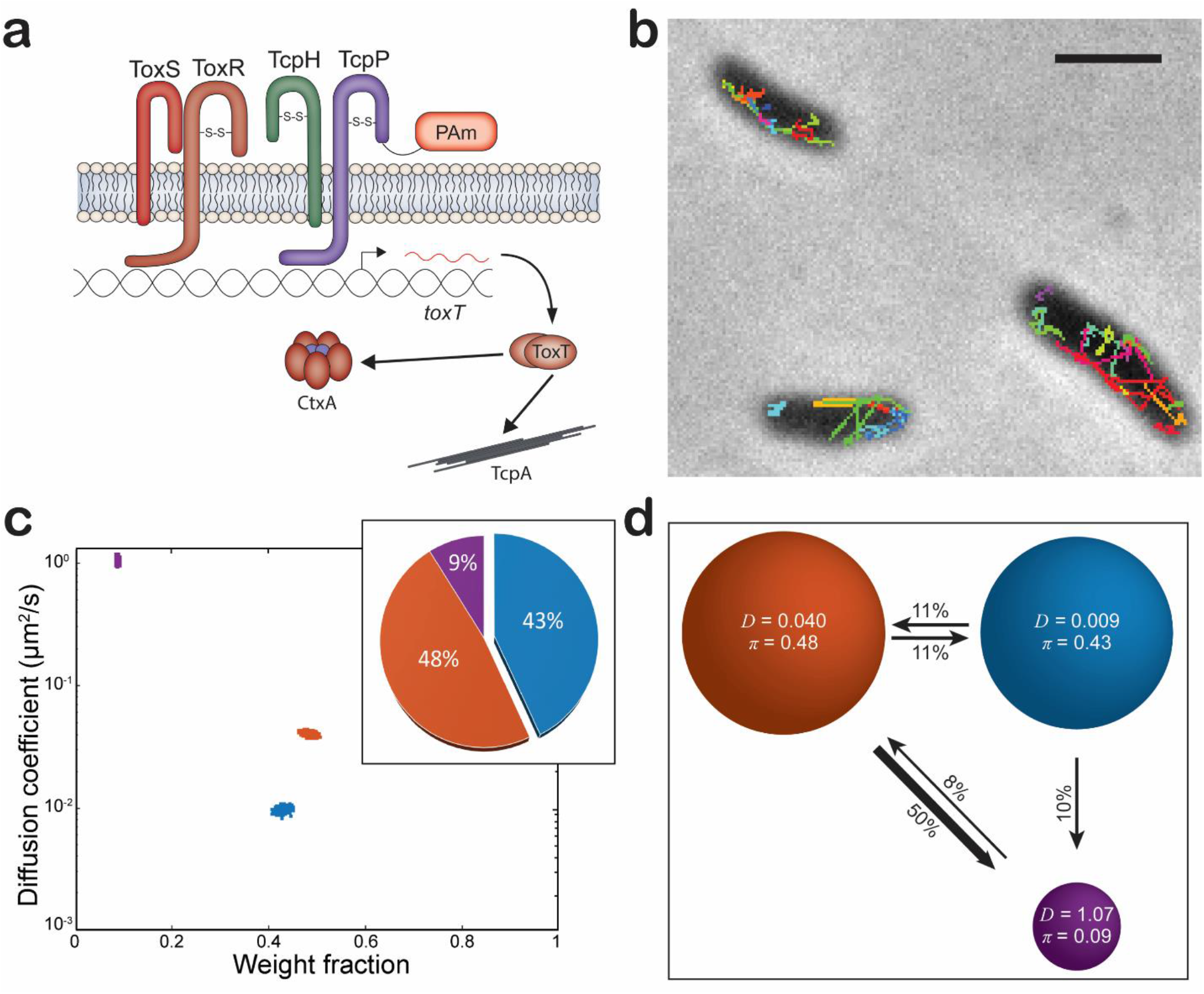
a) Model of *tcpP-PAmCherry*. b) Representative single-molecule trajectory maps overlaid on reverse-contrast bright-field image of *V. cholerae* TcpP-PAmCherry. Only trajectories lasting 0.20 s (5 frames) are shown. Trajectories shown in a variety of colors to show diversity of motion observed. Scale bar: 1 µm. c) Average single-molecule diffusion coefficients and weight fraction estimates for TcpP-PAmCherry in live *V. cholerae* cells grown under virulence-inducing conditions. Single-step analysis identifies three distinct diffusion states (fast – purple, intermediate – orange, and slow – blue, respectively). Each point represents the average single-molecule diffusion coefficient vs. weight fraction of TcpP-PAmCherry molecules in each distinct mobility state at each saved iteration of the Bayesian algorithm after convergence. The dataset contains 54,454 steps from 7,601 trajectories. Inset: percentage (weight fraction) of TcpP-PAmCherry in each diffusion state. Colors as in panel. d) Based on the identification of three distinct diffusion states for TcpP-PAmCherry (three circles with colors as in c and with average single-molecule diffusion coefficient, *D*, indicated in μm^2^/s), the average probabilities of transitioning between mobility states at each step are indicated as arrows between those two circles, and the circle areas are proportional to the weight fractions. Low significance transition probabilities less than 4% are not displayed; for instance the probability of TcpP-PAmCherry molecules transitioning from the fast diffusion state to the slow diffusion state is 1%. Numbers above the arrows indicate the probability of transition.

The high transition probability of TcpP-PAmCherry molecules from the intermediate diffusion state to the fast diffusion state (50%) is unexpected, as the fast diffusion state represents the smallest population of TcpP-PAmCherry molecules (9%), with a low probability (8%) of TcpP-PAmCherry molecules transitioning from the fast diffusion state back to the intermediate diffusion state (Figure 1d). While we cannot directly determine how RIP influences the dynamics of TcpP-PAmCherry, the stark difference in the transition probabilities and the populations of TcpP-PAmCherry in the fast and intermediate diffusion states suggests that fast diffusing TcpP-PAmCherry molecules are potentially sensitive to some form of degradation.

Given this baseline for the dynamics of TcpP-PAmCherry, we hypothesize that: 1) the three diffusion states (slow, intermediate, and fast) are features of TcpP-PAmCherry molecules with three biologically distinct roles; 2) the slow diffusion state is occupied by TcpP-PAmCherry molecules interacting with DNA, such as the *toxTpro*; and 3) the intermediate diffusion state is influenced by ToxR. We further explore these three hypotheses with *V. cholerae* mutants below.

### Mutation of the toxTpro Decreases the Slow Diffusion State Occupancy

We hypothesized that the slow TcpP-PAmCherry diffusion state encompasses molecules specifically interacting with DNA at its binding site in the *toxTpro*. The molecular weight of chromosomal DNA (chromosome 1: 2.96 Mbp) is much higher than that of any protein. Thus, binding of TcpP-PAmCherry to this promoter on the chromosome should result in an extremely low apparent diffusion rate. To test our hypothesis, we removed key binding sites for TcpP (−55 to −37) and both ToxR and TcpP (−112 to +1) in the *toxTpro*, generating *tcpP-PAmCherry toxTpro*Δ(−55–+1) and *tcpP-PAmCherry toxTpro*Δ(−112–+1) (Figure 2), both of which resulted in a drastic reduction in TcpA production, similar to that of a *ΔtcpP* mutant (supplemental Figure 1). *toxT* expression was reduced in *tcpP-PAmCherry toxTpro*Δ(−112–+1), but not in *tcpP-PamCherry toxTpro*Δ(−55–+1) (supplemental Figure 3). It is possible that the *toxTpro*Δ(−55–+1) mutation causes TcpP-PAmCherry and ToxR to stimulate expression of a non-functional *toxT* mRNA. Regardless, loss of either region of the *toxTpro* results in loss of production of the TcpA virulence factor.

**Figure 2:**
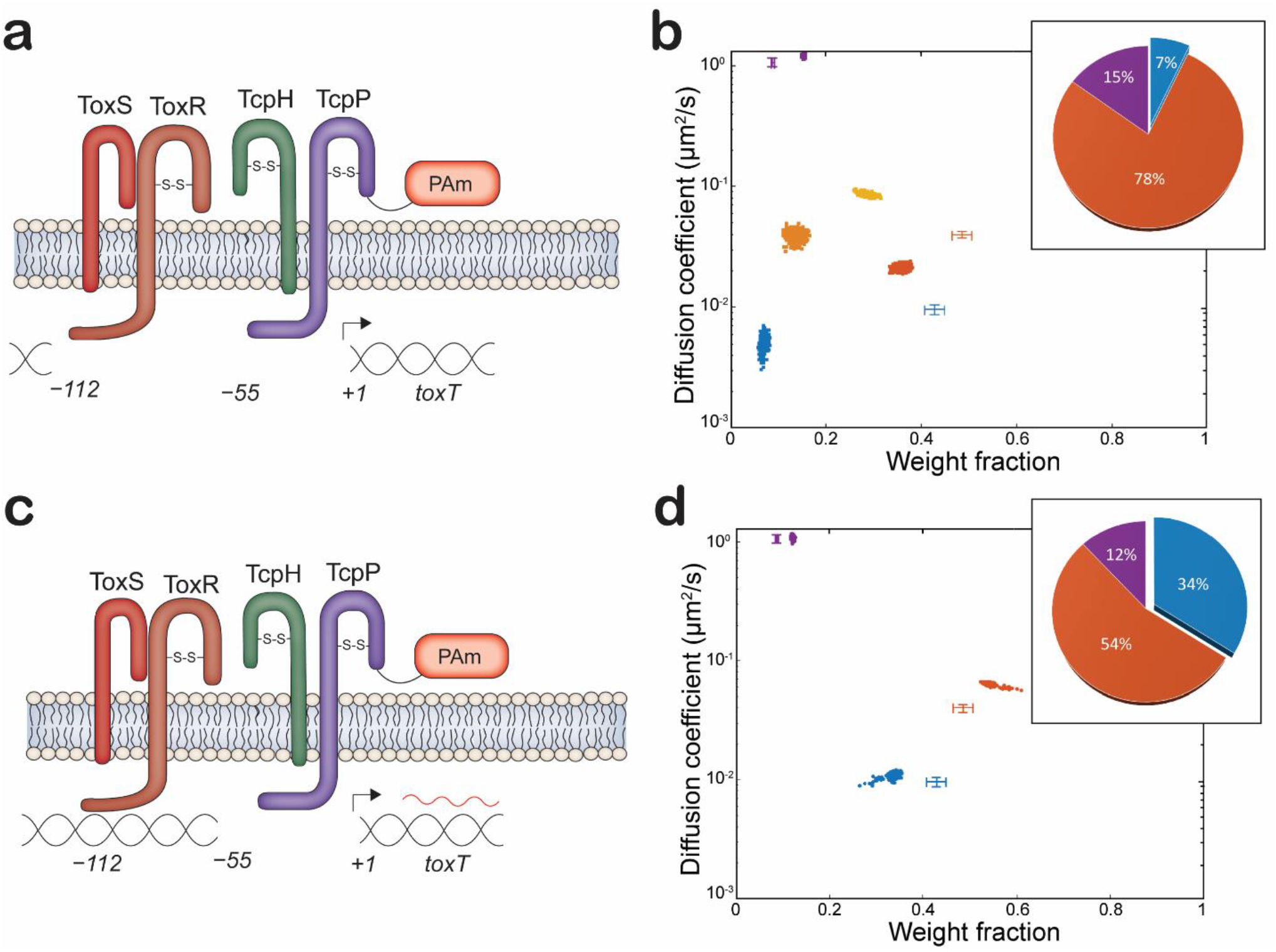
TcpP-PAmCherry diffusion dynamics within live *V. cholerae* cells containing mutated regions of the *toxT* promoter (*toxTpro*). a) and c) Model of *toxTpro* mutations in *tcpP-PAmCherry toxTpro*Δ(−112–+1) and *tcpP-PAmCherry toxTpro*Δ(−55–+1), respectively. b) and d) Average single-molecule diffusion coefficients and weight fraction estimates for TcpP-PAmCherry in live *V. cholerae tcpP-PAmCherry toxTpro*Δ(−112–+1) (b) and *V. cholerae tcpP-PAmCherry toxTpro*Δ(−55–+1) (d) grown under virulence-inducing conditions. Single-step analysis identifies five and three distinct diffusion states (fast – purple, intermediate – orange, light orange, and yellow, and slow – blue, respectively). Each point represents the average single-molecule diffusion coefficient vs. weight fraction of TcpP-PAmCherry molecules in each distinct mobility state at each saved iteration of the Bayesian algorithm after convergence. The dataset contains 104,341 steps from 21,274 trajectories for b and 75,841 steps from 11,624 trajectories for d. The data for TcpP-PAmCherry diffusion in wild type *V. cholerae* cells (Figure 1c) are provided for reference (cross hairs). Insets: Percentage (weight fraction) of TcpP-PAmCherry in each diffusion state. Colors as in panel.

Relative to the wild type (Figure 1), deleting both the ToxR and TcpP binding sites (*toxTpro*Δ(−112–+1)) reduces the percentage of slow diffusing TcpP-PAmCherry to very low levels (7%; Figure 2b). Thus, TcpP-PAmCherry in the slow diffusion state requires *toxTpro*; therefore, we propose molecules in this state are bound to *toxTpro*. On the other hand, loss of the TcpP binding site alone (*toxTpro*Δ(−55–+1)) reduces the percentage of slow TcpP-PAmCherry molecules only subtly (from 43% to 34%; Figure 2d). This result is consistent with earlier observations demonstrating that association with ToxR can restore the function of TcpP variants otherwise unable to bind the *toxTpro* ^18,24^.

Furthermore, our single-step analysis of TcpP-PAmCherry in the *toxTpro*Δ(−112–+1) cells indicates five distinct TcpP-PAmCherry diffusion states, an increase from three states in the wild type (Figure 2b). In particular, the percentage of TcpP-PAmCherry molecules within the intermediate state overall increased (48% to 78%), but our analysis showed that these moderate moving molecules actually cluster into three distinct sub-states (yellow, light orange, and orange, in Figure 2b). These intermediate TcpP-PAmCherry diffusion sub-states appear when TcpP-PAmCherry is unable to associate with the *toxTpro*. Though large-scale changes in the chromosome structure following the promoter deletion may play a role, these intermediate TcpP-PAmCherry diffusion sub-states may represent true biochemical interactions that are too short-lived to precisely distinguish and identify due to our current time resolution of 40 ms/acquisition. Further investigation is required to understand the specific biological roles of these sub-states, but indeed as discussed below, we detect these intermediate sub-states in all the other mutants studied here (Figures 3 and 4).

**Figure 3:**
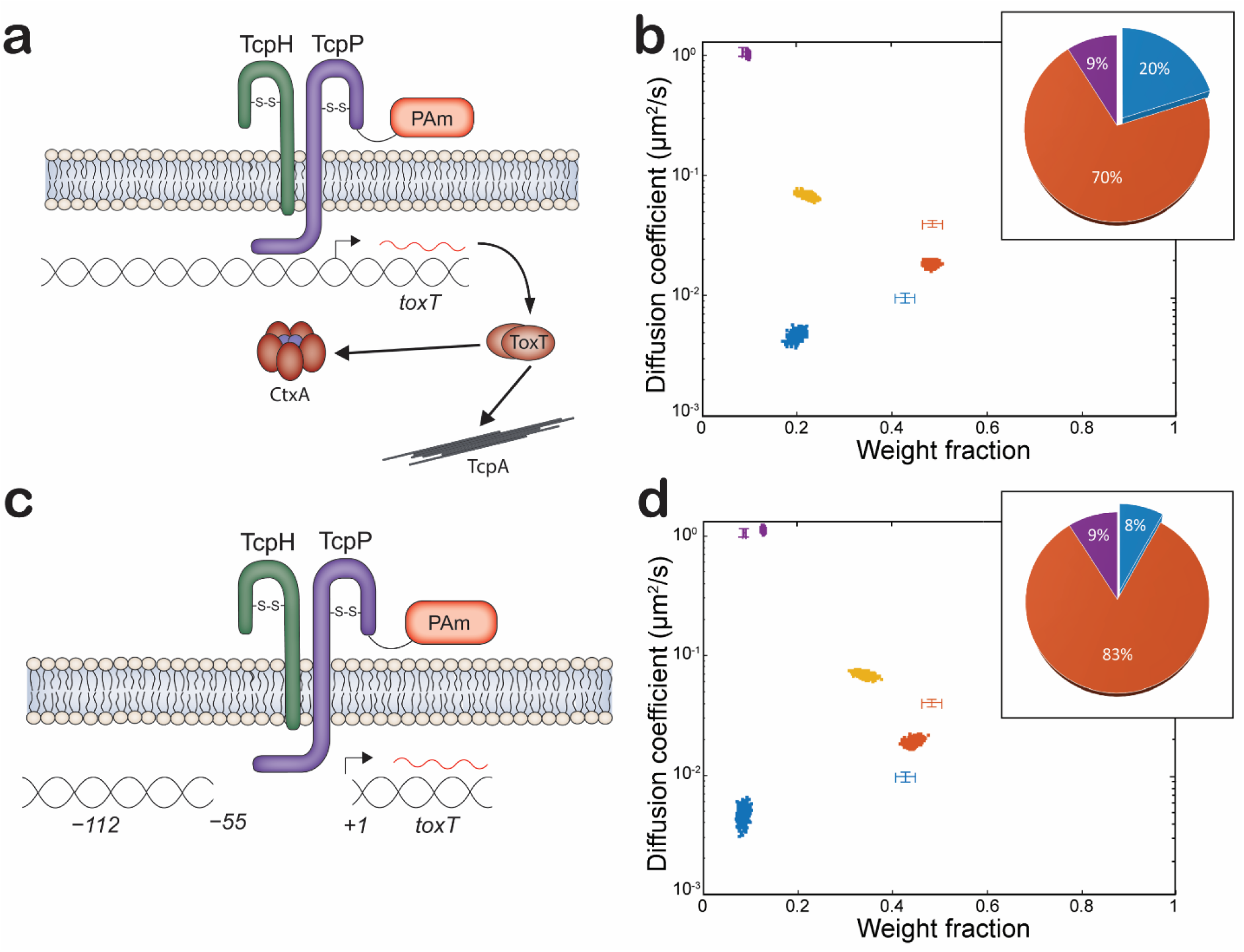
TcpP-PAmCherry diffusion dynamics within live *V. cholerae* cells lacking ToxRS and regions of the *toxT* promoter. a) and c) Model of *tcpP-PAmCherry* Δ*toxRS* and *tcpP-PamCherry* Δ*toxRS toxTpro*Δ(−55–+1), respectively. b) and d) Average single-molecule diffusion coefficients and weight fraction estimates for TcpP-PAmCherry in live *V. cholerae tcpP-PAmCherry* Δ*toxRS* (b) and *V. cholerae tcpP-PAmCherry* Δ*toxRS toxTpro*Δ(−55–+1) (d) grown under virulence-inducing conditions. Single-step analysis identifies four distinct diffusion states (fast – purple, intermediate – yellow and orange, and slow – blue, respectively). Each point represents the average single-molecule diffusion coefficient vs. weight fraction of TcpP-PAmCherry molecules in each distinct mobility state at each saved iteration of the Bayesian algorithm after convergence. The dataset contains 80,005 steps from 11,069 trajectories for b and 58,577 steps from 11,314 trajectories for d. The data for TcpP-PAmCherry diffusion in wild type *V. cholerae* cells (Figure 1c) are provided for reference (cross hairs). Inset: Percentage (weight fraction) of TcpP-PAmCherry in each diffusion state. Colors as in panel.

**Figure 4:**
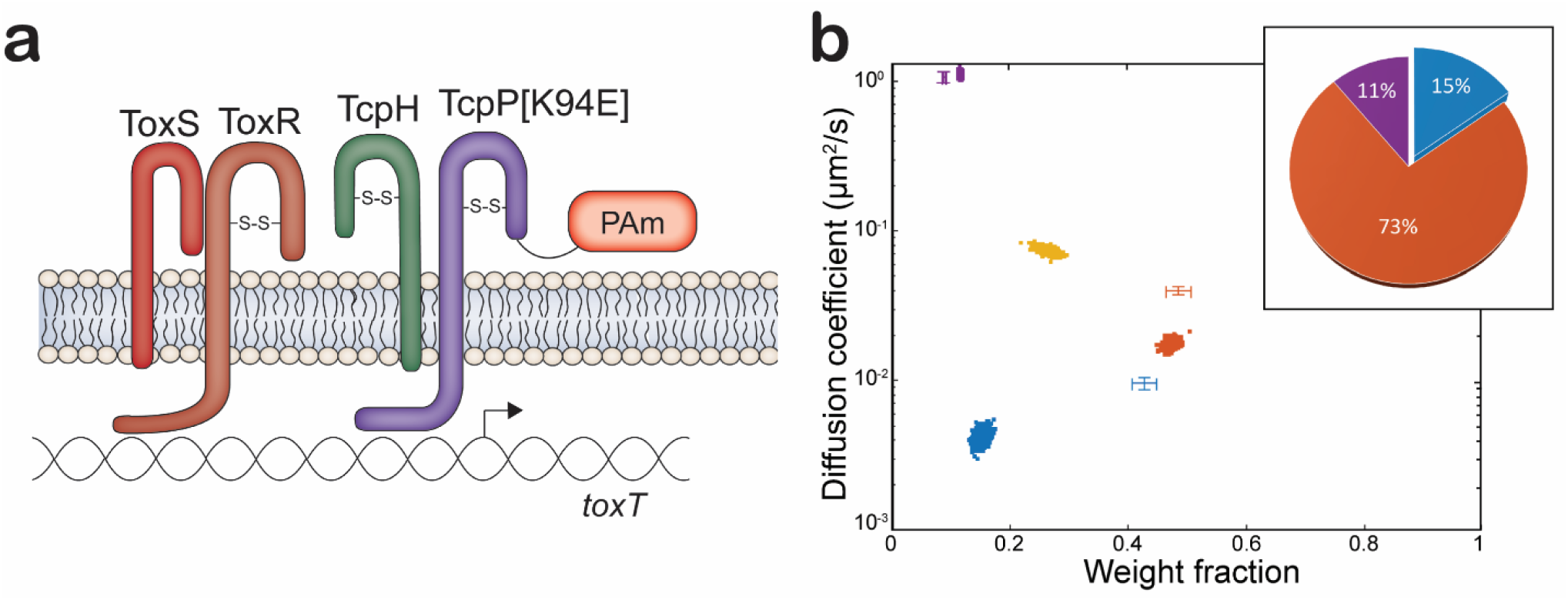
a) Model of *tcpP-[K94E]-PAmCherry*. b) Diffusion dynamics of a DNA binding deficient TcpP-PAmCherry variant within live *V. cholerae* cells. Average single-molecule diffusion coefficients and weight fraction estimates for TcpP-[K94E]-PAmCherry in live *V. cholerae tcpP-[K94E]-PAmCherry* grown under virulence-inducing conditions. Single-step analysis identifies four distinct diffusion states (fast – purple, intermediate – yellow and orange, and slow – blue, respectively). Each point represents the average single-molecule diffusion coefficient vs. weight fraction of TcpP-[K94E]-PAmCherry molecules in each distinct mobility state at each saved iteration of the Bayesian algorithm after convergence. The dataset contains 52,565 steps from 8,056 trajectories. The data for TcpP-PAmCherry diffusion in wild type *V. cholerae* cells (Figure 1c) are provided for reference (cross hairs). Inset: Percentage (weight fraction) of TcpP-[K94E]-PAmCherry in each diffusion state. Colors as in panel.

### ToxR Promotes TcpP-PAmCherry Association with the Slow and Fast Diffusion States

ToxR is a critical regulator of *toxT* expression through its role supporting TcpP interaction with the *toxTpro* ^18,24,25^. Prior studies have shown that TcpP and ToxR interact in response to low oxygen concentrations, and ToxR antagonizes H-NS from the *toxTpro* ^24,48,49^. Several models for TcpP-mediated *toxT* transcription implicate ToxR in recruitment of TcpP molecules to the *toxTpro* ^18,23–25,28,30^. Another model invokes “promoter alteration” to suggest that ToxR promotes TcpP-*toxTpro* interaction by displacing the histone-like protein (H-NS) and altering DNA topology rather than recruiting TcpP molecules to the *toxTpro* ^*28*^.

To examine the role of ToxR in the motion and localization of TcpP-PAmCherry, we deleted *toxR*, and its accessory protein *toxS*, in both the *tcpP-PAmCherry* and the *tcpP-PamCherry toxTpro*Δ(−55–+1) backgrounds, resulting in *tcpP-PAmCherry ΔtoxRS* and *tcpP-PAmCherry ΔtoxRS toxTpro*Δ(−55–+1) genotypes. We found that *tcpP-PAmCherry ΔtoxRS* and *tcpP-PAmCherry ΔtoxRS toxTpro*Δ(−55–+1) cells could activate *toxT* transcription, but only *tcpP-PAmCherry ΔtoxRS* supported virulence factor production (supplemental Figures 1, 3, and 5). Complementation of *tcpP-PAmCherry ΔtoxRS* with *toxR* did not change overall levels of TcpA (supplemental Figure 6). Complementation of *tcpP-PAmCherry ΔtoxRS toxTpro*Δ(−55–+1) with ToxR did not restore TcpA to WT levels (supplemental Figure 6). These data show that TcpP-PAmCherry can stimulate *toxT* expression and bind to the *toxTpro* independent of ToxR. WT TcpP can stimulate *toxT* expression independent of ToxR, but only upon TcpP overexpression ^18,24^. Due to reduced sensitivity of TcpP-PAmCherry to RIP, we measure higher levels of TcpP-PAmCherry relative to TcpP (supplemental Figure 1). This observation suggests that cooperativity between ToxR and TcpP is only necessary when levels of TcpP are low (i.e., when TcpP is sensitive to RIP).

The percentage of slowly-diffusing TcpP-PAmCherry molecules depends on *toxRS*, as deleting *toxRS* reduces this population in *tcpP-PAmCherry ΔtoxRS* from 43% to 20% (Figure 3b). This *toxRS* dependence is maintained even in the absence of the TcpP binding site within the *toxT* promoter; the slow population in *tcpP-PAmCherry ΔtoxRS toxTproΔ*(−55–+1) is reduced to 8% from 34% in *tcpP-PAmCherry toxTpro*Δ(−55–+1) (Figure 3d). Indeed, the TcpP-PAmCherry dynamics are very similar for *tcpP-PAmCherry toxTpro*Δ(−112–+1) (Figure 2b) and *tcpP-PAmCherry ΔtoxRS toxTpro*Δ(−55–+1) (Figure 3d). The major difference between TcpP-PAmCherry diffusion dynamics is the loss of the light orange intermediate diffusion sub-state in *tcpP-PAmCherry ΔtoxRS toxTpro*Δ(−55–+1) (Figure 3d). These data indicate that, in addition to the slow diffusion state, the presence of ToxR is critical for TcpP-PAmCherry molecules to exist in one of the intermediate sub-state diffusion states (i.e., the light orange diffusion state).

As shown in Figure 1d, we found that TcpP-PAmCherry molecules do not freely transition between all the diffusion states: the intermediate diffusion state is an important diffusion state for TcpP-PAmCherry molecules to transition between the fast and the slow diffusion states. Since the ToxR-TcpP interaction is proposed to enable TcpP to associate with the transcription complex at *toxTpro* ^18,24^, we reasoned that ToxR is responsible for the preferred intermediate-to-slow state transition of TcpP-PAmCherry. However, in Δ*toxRS* (Figure 3b) like in the wild-type (Figure 1c), only TcpP-PAmCherry molecules in the slowest of the intermediate diffusion sub-states were likely to transition to the slow diffusion state (orange and blue diffusion states, respectively, supplemental Figure 7). These transition probabilities suggest that ToxR is not responsible for the restricted transition of TcpP-PAmCherry between the slow and fast diffusion states. Furthermore, the absence of ToxR reduced the probability of TcpP-PAmCherry entering the fast diffusion state and increased the probability of TcpP-PAmCherry leaving the fast diffusion state (Figure 1d and supplemental Figure 7b). Taken together, these data indicated that ToxR sequesters a portion of the total TcpP-PAmCherry population away from the *toxTpro*. We reasoned that increased levels of ToxR might sequester TcpP molecules to an inactive state (represented by the intermediate diffusion state). To test this hypothesis, we overexpressed ToxR in a *tcpP-PAmCherry* background and quantified virulence factor expression (i.e., TcpA) (supplemental Figure 8). We found that elevated ToxR levels reduced virulence factor levels in both WT and *tcpP-PAmCherry* cells.

### Mutation of the TcpP Helix-Turn-Helix Domain Reduces the Percentage of Slowly Diffusing TcpP-PAmCherry

Based on results shown in Figure 1c, we proposed that TcpP-PAmCherry molecules in the slow diffusion state are bound to *toxTpro*, and we found that removing the *toxTpro* binding sites (Figure 2) or eliminating *toxR* (Figure 3) significantly reduces this bound state population. Previous studies demonstrated that TcpP does not require DNA binding capability to activate *toxT* expression if ToxR is present ^18,24^. To examine this finding further by SMT, we used a *tcpP-PAMCherry* allele with a mutation (K94E) that inhibits TcpP from binding to the *toxTpro* ^*24*^. This mutation results in greatly reduced *toxT expression* and TcpA levels (supplemental Figures 1 and 3). The levels of TcpP[K94E]-PAmCherry is elevated compared with TcpP-PAmCherry (supplemental Figure 1), consistent with earlier evidence that the K94E substitution increases TcpP stability ^24^. In addition to TcpP[K94E]-PAmCherry being unable to stimulate *toxT* expression, a lower percentage of TcpP[K94E]-PAmCherry molecules are detected in the slowest-diffusing state than for TcpP-PAmCherry (15% vs. 43%; Figure 4b). Furthermore, TcpP[K94E]-PAmCherry molecules have an additional intermediate diffusion sub-state, similar to both *tcpP-PamCherry ΔtoxRS* and *tcpP-PAmCherry ΔtoxRS toxTproΔ*(−55–+1) (Figure 4b).

## Discussion

How MLTAs find their target sequences from the membrane represents a major gap in knowledge. Here, we started to address this by investigating single-molecule dynamics of TcpP-PAmCherry. Taken together with previous work, the data presented here demonstrate that TcpP-PAmCherry molecules diffuse in at least three distinct biophysical states (fast, intermediate, and slow diffusion), but do not freely transition between all diffusion states ^44^. We hypothesized that each of these biochemical states have distinct biological roles. Specifically, we hypothesized that the slow diffusion state represented TcpP-PAmCherry molecules interacting with the *toxTpro*. To test this hypothesis, we made targeted deletions to the *toxTpro* and of *toxRS*, and we mutated the TcpP DNA binding domain (K94E). Our biophysical measurements of these mutations support the hypothesis that the slow diffusion state is occupied by TcpP-PAmCherry molecules interacting specifically with DNA at *toxTpro*. Additionally, we observed that TcpP-PAmCherry molecules transition to the slow diffusion state from the intermediate diffusion state prior to transitioning, and that ToxR is not responsible for this transition specificity. These data support a modified promoter alteration model ^28^ in which ToxR binds to the distal region of the *toxTpro* to promote TcpP binding to the proximal region of the *toxTpro* or, in the absence of its binding site, ToxR directly interacts with TcpP to stimulate *toxT* expression. Our data do not suggest that ToxR directs or recruits TcpP to the *toxTpro*.

While ToxR is critical for TcpP to stimulate *toxT* expression^18,24,27^, our data demonstrate that TcpP-PAmCherry can support *toxT* expression and virulence factor production without ToxR, which may be a consequence of the greater stability of TcpP-PAmCherry compared to native TcpP (supplemental Figure 1 and 3). Moreover, our single-molecule imaging finds a higher percentage of the TcpP-PAmCherry molecules in the slow diffusion state in *tcpP-PamCherry ΔtoxRS* cells compared to *tcpP-PAmCherry ΔtoxRS toxTpro*Δ(−55–+1) (Figure 3). In addition, prior DNAse I footprinting experiments have demonstrated that in cells lacking *toxR* TcpP protects a larger region of the *toxTpro* (−100 to −32), i.e., TcpP protects most of the ToxR binding and TcpP binding sites in *ΔtoxRS* ^18^. Taken together, these results indicate that: 1) ToxR is not essential for TcpP to locate the *toxTpro;* and 2) TcpP is able to interact with the *toxTpro* independent of ToxR. These data support the promoter alteration model ^28^ in which ToxR assists TcpP in stimulating *toxT* transcription, but does not necessarily recruit TcpP to the *toxTpro*.

In addition, our data also show that Δ*toxRS* reduces the percentage of DNA-bound TcpP-PAmCherry (Figure 3) and decreases the probability of TcpP-PAmCherry molecules transitioning from the intermediate state to the bound state (7% in Δ*toxRS* vs. 11% in *tcpP-PAmCherry*) (supplemental Figure 7B). Despite this small reduction in the transition probability, TcpP-PAmCherry stimulates WT *toxT* expression independent of ToxR (supplemental Figure 3). The histone-like protein H-NS can bind to the *toxTpro* suppressing *toxT* expression ^50^. ToxR antagonizes H-NS, thereby relieving this repression^48^. TcpP is capable of stimulating *toxT* expression without ToxR, and thereby in the presence of H-NS ^18,24^. However, this requires overexpression of TcpP, which does not support WT levels of *toxT* expression ^18,24^. This suggests that TcpP competes poorly with H-NS for the *toxTpro* relative to ToxR. We hypothesize that the small reduction in transition probability of TcpP-PAmCherry molecules to the slow diffusion state upon deletion of *toxRS* is due to increased crowding of the *toxTpro* by H-NS. Additional experiments are needed to test this hypothesis. Deletion of both *toxRS and toxTpro*(−55–+1) (i.e., the TcpP binding site) restores the probability of TcpP-PAmCherry molecules transitioning from the intermediate state to the bound state (back to 12% vs. 11% in cells wild type for those *tcpP-PAmCherry*) (supplemental Figure 7c). It is unclear why deletion of the TcpP binding site in the *toxTpro* would restore the probability of TcpP-PAmCherry molecules transitioning to the bound state. However, these data also support an alternative model in which, rather than ToxR recruiting TcpP to the *toxTpro*, ToxR assists TcpP to stimulate *toxT* transcription once TcpP associates with the *toxTpro*.

Under certain conditions ToxR can negatively influence *toxT* expression. In response to stationary-phase accumulation of the cyclic di-peptide cyclic phenylalanine-proline (cyc-phe-pro), ToxR stimulates production of LeuO, resulting in down-regulation of the *tcpP* regulator *aphA* ^51,52^. Our data suggests that ToxR can also reduce *toxT* expression by influencing TcpP-PAmCherry single molecule dynamics (supplemental Figure 7b). Deletion of *toxRS* reduces the overall probability of TcpP-PAmCherry molecules transitioning between the intermediate and fast diffusion states (supplemental Figure 7b). Moreover, elevated levels of ToxR reduce virulence factor production (supplemental Figure 8), suggesting that ToxR can antagonize virulence factor production by promoting transition of TcpP molecules to the fast diffusion state. A similar phenotype has been reported previously ^18^. Lastly, prior electrophoretic mobility shift assays also indicate that ToxR can sequester TcpP from the *toxTpro*. In *ΔtoxRS* cells TcpP is able to bind to the *toxTpro* -73_+45 (*toxTpro* lacking the ToxR binding region), but not in the presence of ToxR molecules ^18^. It remains unclear how ToxR promotes transition of TcpP-PAmCherry molecules to the fast diffusion state. However, we hypothesize that ToxR promotes transition of TcpP molecules to the fast diffusion state to prevent aberrant *toxT* expression. Our data also suggest that TcpP-PAmCherry molecules in the fast diffusion state may be more sensitive to RIP than molecules in the intermediate and slow diffusion states. Thus, ToxR mediated transition of TcpP molecules to the fast diffusion state may promote turnover of TcpP molecules via RIP. Follow-up experiments are required to test this hypothesis.

Currently, the biological roles of the intermediate diffusion states (or intermediate diffusion sub-states) are unclear, but the intermediate states are certainly important, as TcpP molecules transition to the *toxTpro*-bound state from them. There is nearly a 10-fold difference in diffusion coefficients between the slow and intermediate diffusion states (0.044 µm^2^/sec vs. 0.006 µm^2^/sec respectively; Figure 1c). This difference cannot be explained by dimerization or interaction of ToxR and TcpP-PAmCherry alone: the mobility of membrane-localized proteins scales linearly with the number of transmembrane helices, such that increasing the number of transmembrane helices via dimerization from one to two would only reduce the diffusion coefficient by a factor of two ^53^. One possibility is that TcpP-PAmCherry molecules undergo fast diffusion in less protein dense areas of the cytoplasmic membrane relative to TcpP-PAmCherry molecules undergoing intermediate diffusion. Alternatively, it is possible that the diffusion coefficients of TcpP-PAmCherry molecules in the intermediate state are undergoing non-specific interactions with DNA whereas the slowest TcpP-PAmCherry molecules are specifically bound at *toxTpro*. Our data show that there are some slow moving TcpP-PAmCherry molecules when major regions of the *toxTpro* are deleted or when key residues within the DNA binding domain of TcpP are mutated (i.e., *tcpP[K94E]-PAmCherry;* Figure 2 and 4). When considering our alternative model of non-specific DNA binding by TcpP, these data suggest two possibilities: 1) TcpP-PAmCherry molecules in the slow diffusion state represent TcpP molecules that make both specific and non-specific interactions with DNA; or 2) TcpP-PAmCherry molecules in the slow diffusion state interact specifically with non-*toxTpro* DNA (i.e., TcpP regulates additional genes). Several genes appear to have altered gene expression upon deletion of *tcpPH* ^54^. However, these experiments have yet to be replicated. Thus, future experiments would be required to test these hypotheses.

These results provide deep insights that further expand the model of cooperativity between ToxR and TcpP-PAmCherry. Our data demonstrate that ToxR assists TcpP to associate with the *toxTpro* even in the absence of the TcpP binding site, further supporting the established model of cooperativity between TcpP and ToxR. The data also show that TcpP can locate the *toxTpro*, interact with the *toxTpro*, and stimulate *toxT* expression independent of ToxR. This supports the promoter alteration model in which TcpP molecules independently associate with the *toxTpro* while ToxR enhances this association by displacing H-NS and altering *toxTpro* topology to stimulate *toxT* transcription. Furthermore, these data show that ToxR promotes transition of TcpP molecules to the fast diffusion state, shifting the equilibrium of TcpP molecules away from the *toxTpro*. The mechanism and the biological significance of ToxR promoting transition of TcpP molecules to the fast diffusion state is currently unclear but will be the subject of future investigation. Lastly, since MLTAs are found in both Gram-negative and Gram-positive bacteria, targeting MLTAs could be an effective strategy to treat bacterial infections without exacerbating the global antibiotic resistance crisis.

## Author Contributions

Study design: Lucas Demey, Josh Karslake, Victor DiRita, and Julie Biteen. Microscopy and data analysis: Anna Calkins, Josh Karslake, Eric Donarski. Strain construction: Lucas Demey. Biochemical assays: Lucas Demey. Drafting of the manuscript: Lucas Demey and Anna Calkins. Critical revision of the manuscript: Anna Calkins, Lucas Demey, Josh Karslake, Julie Biteen, and Victor DiRita.

## Data availability

The data presented here will be made available from the corresponding authors upon request.

## Acknowledgements

This work was supported by the National Institutes of Health (grant R21-GM128022) to J.S.B., and the Rudolph Hugh Endowment at Michigan State University (V.J.D.). L.M.D. was a trainee of the Institutional Pharmacological Sciences Training Program at MSU (2T32 GM092715).

## Experimental Procedures

### Bacterial strains and growth conditions

*Escherichia coli* and *V. cholerae* strains used here can be found in supplemental Table 1. Unless otherwise stated, *E. coli* and *V. cholerae* cells were grown on Lysogeny Broth (LB) plates, or in LB broth at 210 rpm, at 37 °C. LB was prepared according to previous descriptions ^55^. To stimulate virulence, *V. cholerae* cells were diluted from overnight cultures in LB broth and subcultured into virulence-inducing conditions: (LB pH 6.5, 110 rpm, 30 °C; filter sterilized). Here, the LB pH was adjusted by adding HCl (1 N) to pH 6.5 (+/-0.05) and then the media was filter-sterilized to maintain pH. Where appropriate, antibiotics and cell wall intermediates were added at the following concentrations: streptomycin (100 µg ml^−1^), ampicillin (100 µg ml^−1^), and diaminopimelic acid (DAP) (300 µM).

### Plasmid construction

Plasmid vectors were purified using the Qiagen mini prep kit. Plasmid inserts were amplified from *V. cholerae* genomic DNA using Phusion high-fidelity polymerase (Thermo Scientific). Splicing by overlap extension was used to combine the entire plasmid insert sequences together, see Supplemental Table 2 for primer list. Plasmid vector was digested by restriction digestion using KpnI-HiFi and XbaI (New England BioLabs) at 37 °C for 2 hrs. After digestion the plasmid vector and insert were added to Gibson assembly master mix (1.5 µl insert, 0.5 µl vector, 2 µl master mix) (New England BioLabs) and incubated at 50 °C for 1 hr. Assembled plasmid was electroporated into *E. coli* λpir cells and recovered on LB plates with ampicillin and DAP.

### Bacterial strain construction

Strain construction follows the protocol outlined in reference ^56^. Briefly, *E. coli* λpir harboring the pKAS plasmid and the donor *V. cholerae* strain were incubated in LB (broth or agar) supplemented with DAP overnight at 37 °C. The remaining cells were then spread on LB plates containing ampicillin or TCBS plates containing ampicillin. Counter selection for loss of the pKAS construct by *V. cholerae* cells was done by incubating cells in LB broth for 2 hrs and then for 2 hrs with 2500 µg ml^−1^ streptomycin (both at 37 °C, 210 rpm). 20 µl of this culture was spread onto LB plates containing 2500 µg ml^−1^ of streptomycin and incubated overnight at 37 °C. Streptomycin-resistant colonies were screened for the chromosomal mutation of interest via colony polymerase chain reaction (PCR) using Taq DNA Polymerases (Thermo Fisher). Genomic DNA was purified from possible mutants and sequenced (Genewiz) to validate the exchange. Because *tcpP* and *tcpH* are encoded by on overlapping open reading frames, *tcpH* was cloned downstream of PAmCherry to maintain its expression, and a stop codon was introduced within the first three codons of the native *tcpH* coding sequence to prevent out-of-frame translation of PAmCherry.

### Growth Curves

*V. cholerae* strains were initially grown on LB plates containing streptomycin (100 µg ml^−1^) overnight at 37 °C, then an individual colony was picked and grown overnight in LB broth at 37 °C. *V. cholerae* cells were diluted to an optical density (OD_600_) of 0.01 from the overnight LB broth into a 96 well plate (Cell Pro) with 200 ul of virulence-inducing media per well. The plate was then incubated at 30 °C with shaking every 30 min before each measurement in a SPECTROstar Omega plate reader (BMG LABTECH).

### Real-time quantitative PCR (RT-qPCR)

RNA was extracted from *V. cholerae* cells grown under virulence-inducing conditions. RNA was preserved by resuspending pellet cells in 1 ml Trizol (Sigma aldrich) and then purified using an RNeasy kit (Qiagen). RNA was further purified with Turbo DNase treatment. RNA quantity and quality were measured via UV-Vis spectrophotometry (NanoDrop ND-1000) and by detection of large and small ribosomal subunits via 2% agarose gel. RNA was then converted to cDNA using Superscript III reverse transcriptase (Thermo Scientific). RT-qPCR was performed using 5 ng of cDNA in SYBR green master mix (Applied Biosystems). RecA was used as a housekeeping gene of reference to calculate the threshold values (ΔΔC_T_) ^57^. See supplemental Table 2 for primers.

### Protein electrophoresis and immunodetection

After lysis, total protein concentration samples were measured via Bradford assay. Samples were subsequently diluted to 0.5 µg total protein/µl. All SDS page gels contained 12.5 % acrylamide and were run at 90 – 120 volts for 1.5 hrs. Proteins were transferred to nitrocellulose membranes using a semi-dry electroblotter (Fisher Scientific) overnight at 35 mA or for 2 hrs at 200mA. Membranes were blocked with 5 % non-fat milk, 2 % bovine serum albumin in Tris-buffered saline, 0.5 % Tween-20 (TBST) for 1 hr. Membranes were then incubated with primary antibody (α-TcpA 1:100,000; α-TcpP 1:1,000; α-TcpH 1:500; α-ToxR 1:50,000; α-mCherry 1:1,000) diluted in TBST and non-fat Milk (2.5 % w/vol) for an additional hour at room temp with shaking. Membranes were then washed 3 times with TBST. Secondary antibody (Goat anti-Rabbit IgG-HRP 1:2,000) (Sigma) was diluted in TBST and non-fat milk (2.5 % w/vol). Secondary antibody was incubated with the membranes for an additional hour at room temperature with shaking. Membranes were washed again with TBST 3 times and then incubated with SuperSignal HRP Chemiluminescence substrate (Thermo Fisher). Membranes were imaged with an Amersham Imager 600.

### Single-Molecule Microscopy

*V. cholerae* strains were grown on LB plates containing streptomycin (100 µg ml^−1^) overnight at 37 °C, then an individual colony was picked and grown overnight in LB broth at 37 °C. *V. cholerae* cells were diluted from LB broth into virulence-inducing conditions and grown until they reached mid log-phase. They were then washed and concentrated in M9 minimal media with 0.4 % glycerol. A 1.5 μl droplet of concentrated cells was placed onto an agarose pad (2 % agarose in M9, spread and flattened on a microscope slide) and covered with a coverslip. Cells were imaged at room temperature using an Olympus IX71 inverted epifluorescence microscope with a 100x 1.40 NA oil-immersion objective, a 405-nm laser (Coherent Cube 405-100; 50 W/cm^2^) for photoactivation and a co-aligned 561-nm laser (Coherent-Sapphire 561-50; 210 W/cm^2^) for fluorescence excitation. Fluorescence emission was filtered with appropriate filters and captured on a 512 by 512 pixel Photometrics Evolve EMCCD camera. To prevent higher-order excitation during photoactivation, a pair of Uniblitz shutters controlled the laser beams such that samples were exposed to only one laser at a time. During imaging, the cells were given a 40-ms dose of 405-nm light every 90 s. Images were collected continuously every 40 ms and acquisitions lasted 5 – 7 min each.

## Data Analysis

Recorded single-molecule positions were detected and localized based on point spread function fitting using home-built code, SMALL-LABS ^58^. This program reduces biases due to background subtraction, increasing the precision of each molecule localization. Subsequent localizations of the same molecule were then connected into trajectories using the Hungarian algorithm ^59,60^. All trajectories from each movie for a given condition were combined and analyzed together using the Single-Molecule Analysis by Unsupervised Gibbs sampling (SMAUG) algorithm ^44^. This algorithm considers the collection of steps in all trajectories and uses a Bayesian statistical framework to estimate the parameters of interest: number of mobility states, diffusion coefficient, weight fraction, transition probabilities between states, and noise.

