## Supplemental Data for "Independent Promoter Recognition by TcpP Precedes Cooperative Promoter Activation by TcpP and ToxR"

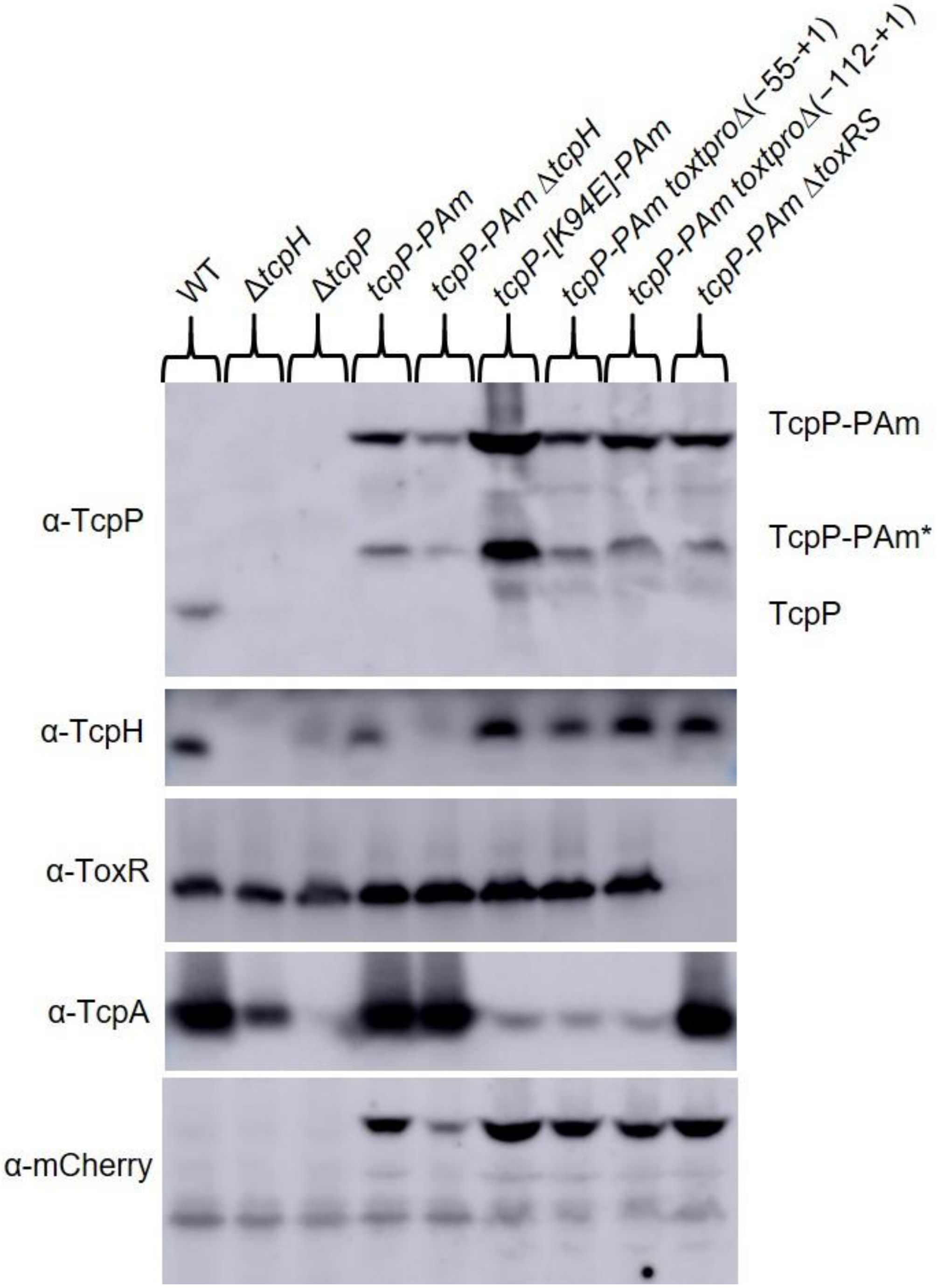


**Supplemental Figure 1:** Western blots of cultures grown under virulence-inducing conditions for 6 hrs, see methods for primary antibody dilution. Photoactivatable mCherry (PAmCherry) is fused to the C-terminus of TcpP and is under the control of its endogenous promoter on the chromosome. Addition of PAmCherry to TcpP results in two species: TcpP-PAmCherry (~70KDa)

and TcpP-PAmCherry* (~36KDa). Deletion of *tcpH* yields lower levels of TcpP-PAmCherry and TcpP-PAmCherry*, likely due to an increase in regulated intramembrane proteolysis (RIP).

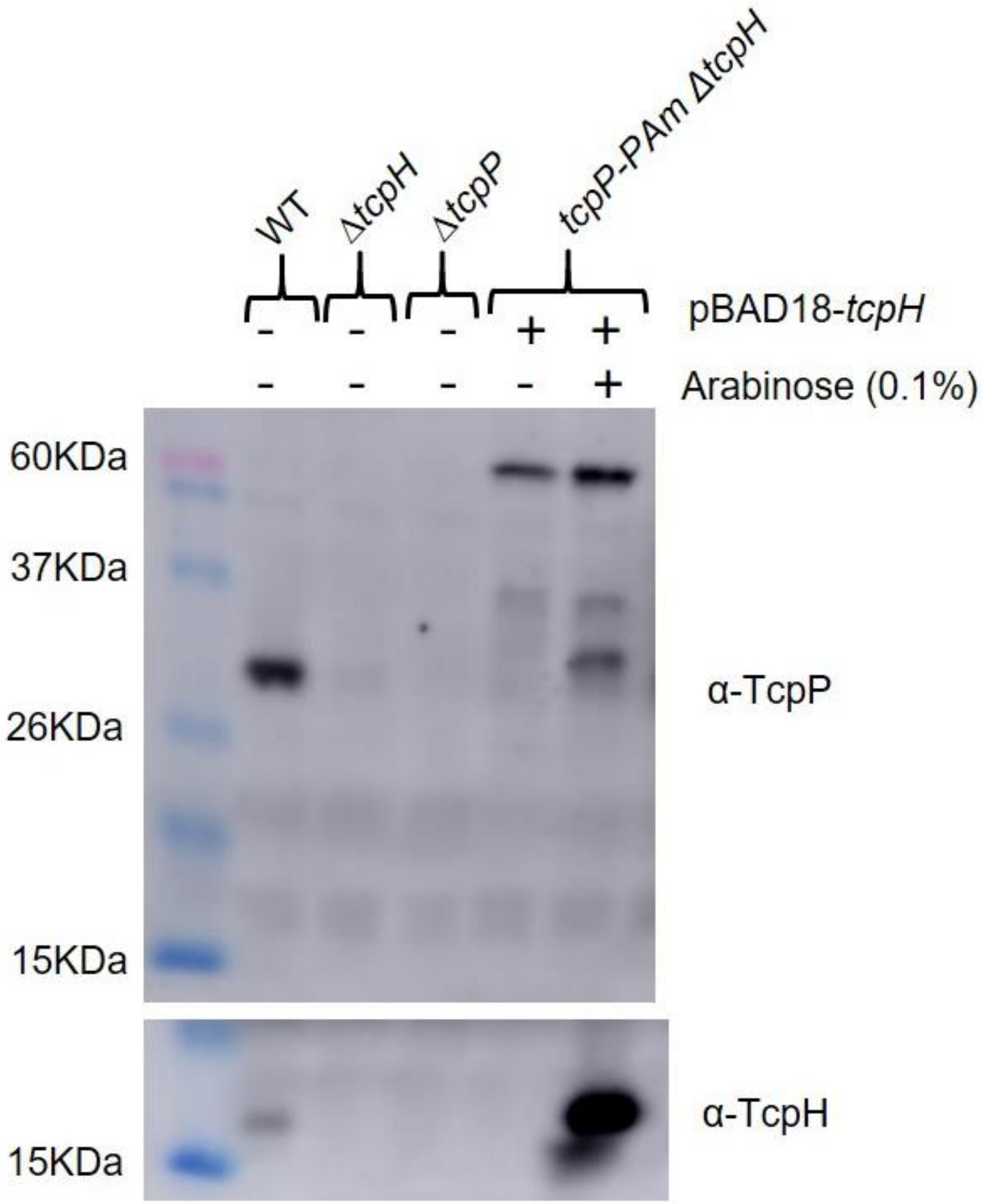


**Supplemental Figure 2:** Western blots of cultures grown under virulence-inducing conditions for 6 hrs with or without arabinose, see methods for primary antibody dilution. *tcpP-PAmCherry* Δ*tcpH* cells harbor an arabinose-inducible vector (pBAD18) encoding *tcpH*. Ectopic expression of *tcpH* complemented deletion of *tcpH*. Complementation of *tpcH* also resulted in an additional TcpP band, ~29KDa, that corresponds to native TcpP.

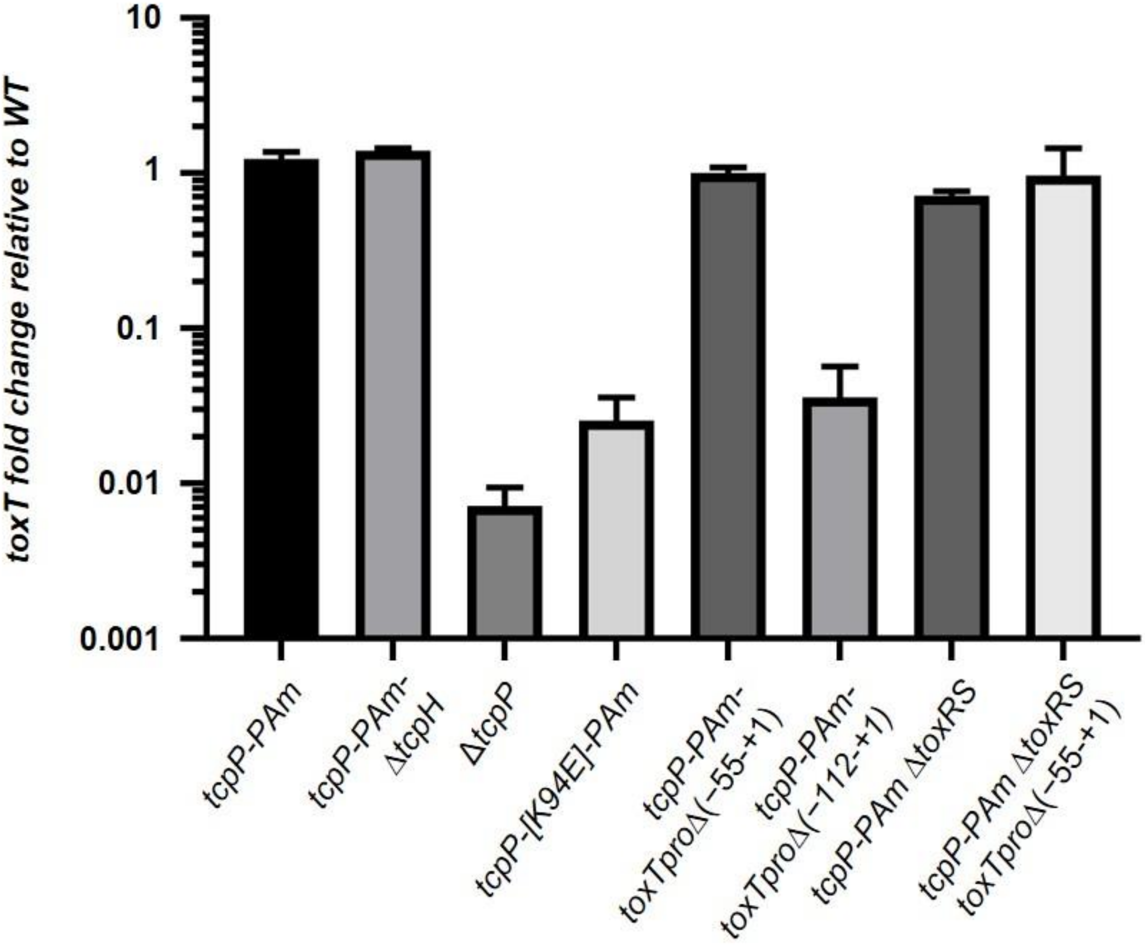


**Supplemental Figure 3:** Average *toxT* fold change, relative to WT, across three biological replicates (determined via the $\Delta\Delta C_T$ method) [61]. mRNA was collected from cells after 2 hrs in virulence-inducing conditions, and error bars represent standard error of the mean.

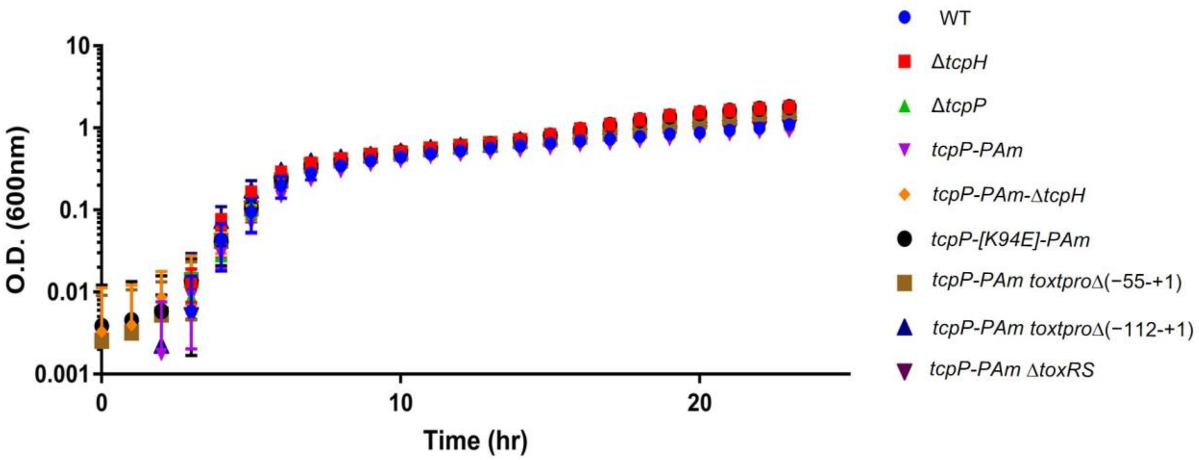


**Supplemental Figure 4:** in vitro growth curve under virulence-inducing conditions. Optical density (O.D.) values are the average of three biological replicates and error bars represent standard error of the mean.

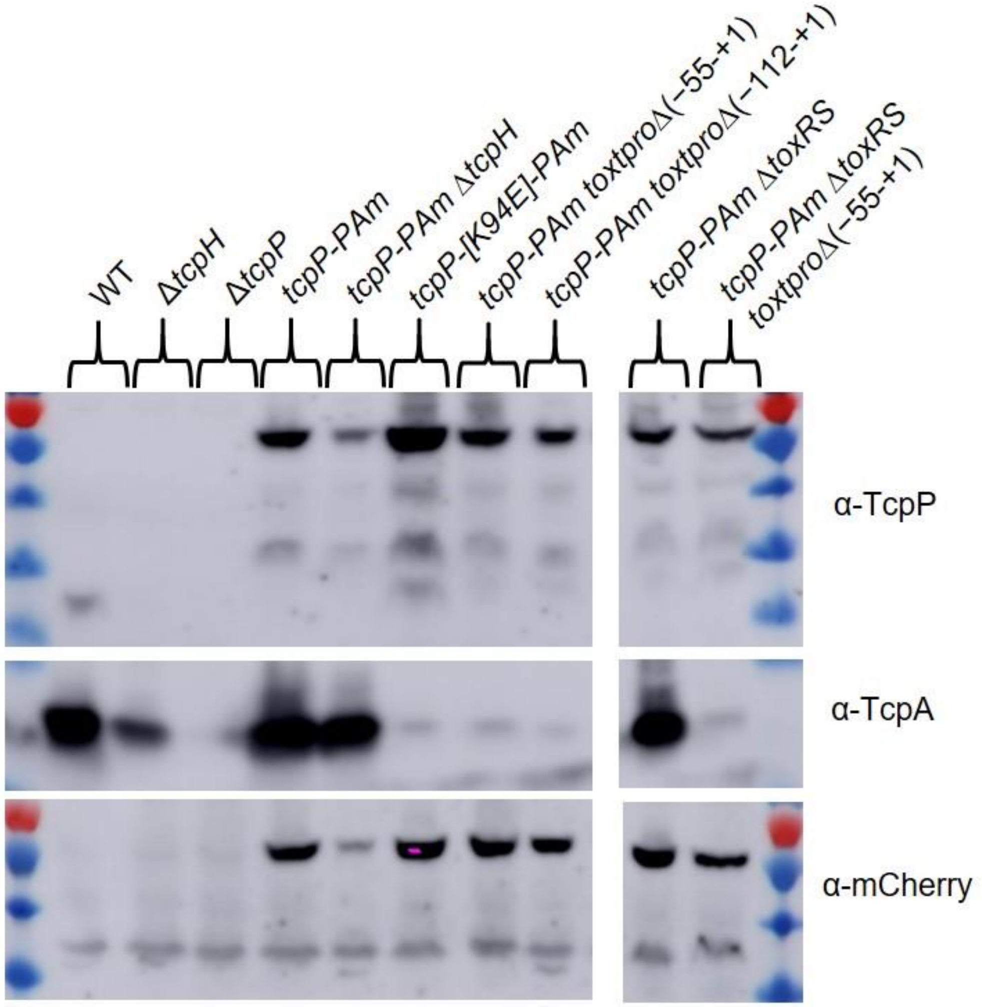


**Supplemental Figure 5:** Western blots of cultures grown under virulence-inducing condition for 6 hrs. See methods for primary antibody dilutions.

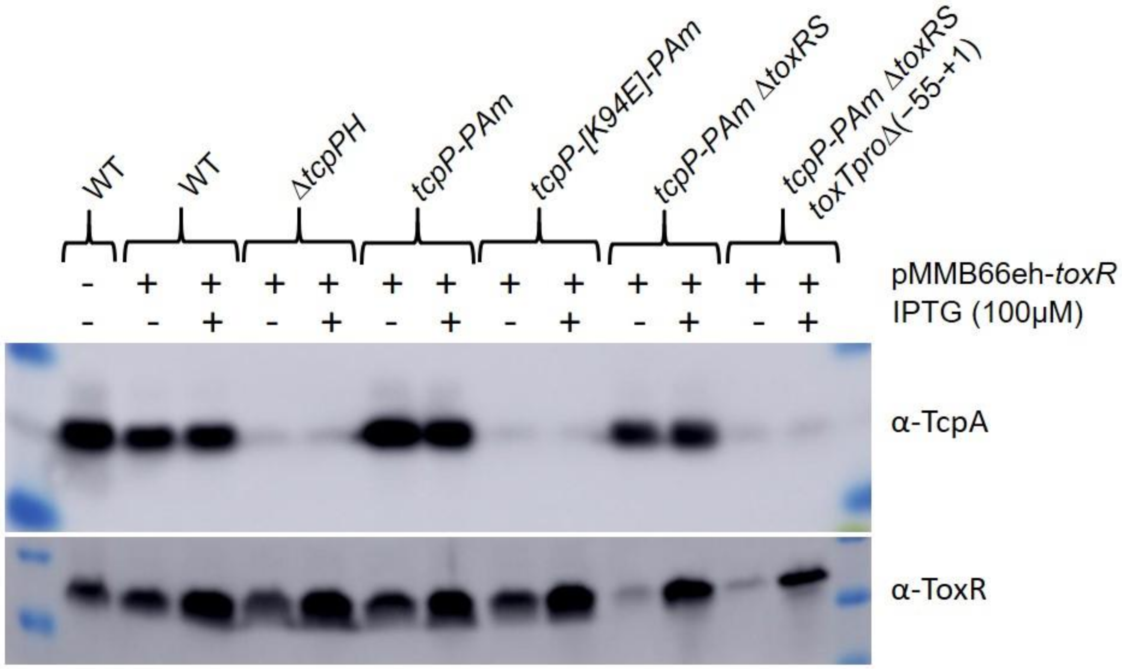


**Supplemental Figure 6:** Complementation and overexpression of ToxR from the pMMB66eh plasmid. Western blots of cellular lysates collected after growth under virulence-inducing conditions for 6 hrs with or without IPTG, see methods for primary antibody dilution. ToxR does not stimulate TcpA production without TcpPH, and ToxR cannot complement TcpPK94E-PAmCherry or *toxTpro*Δ(−55–+1). Low levels of ToxR were detected in *tcpP-PAmCherry* Δ*toxRS* and *tcpP-PAmCherry* Δ*toxRS toxTpro*Δ(−55–+1) without IPTG, likely due to leaky expression of *toxRS* at the IPTG promoter. Multiple copies of the lac promoter are known to result in leaky expression due to insufficient levels of LacI [62,63].

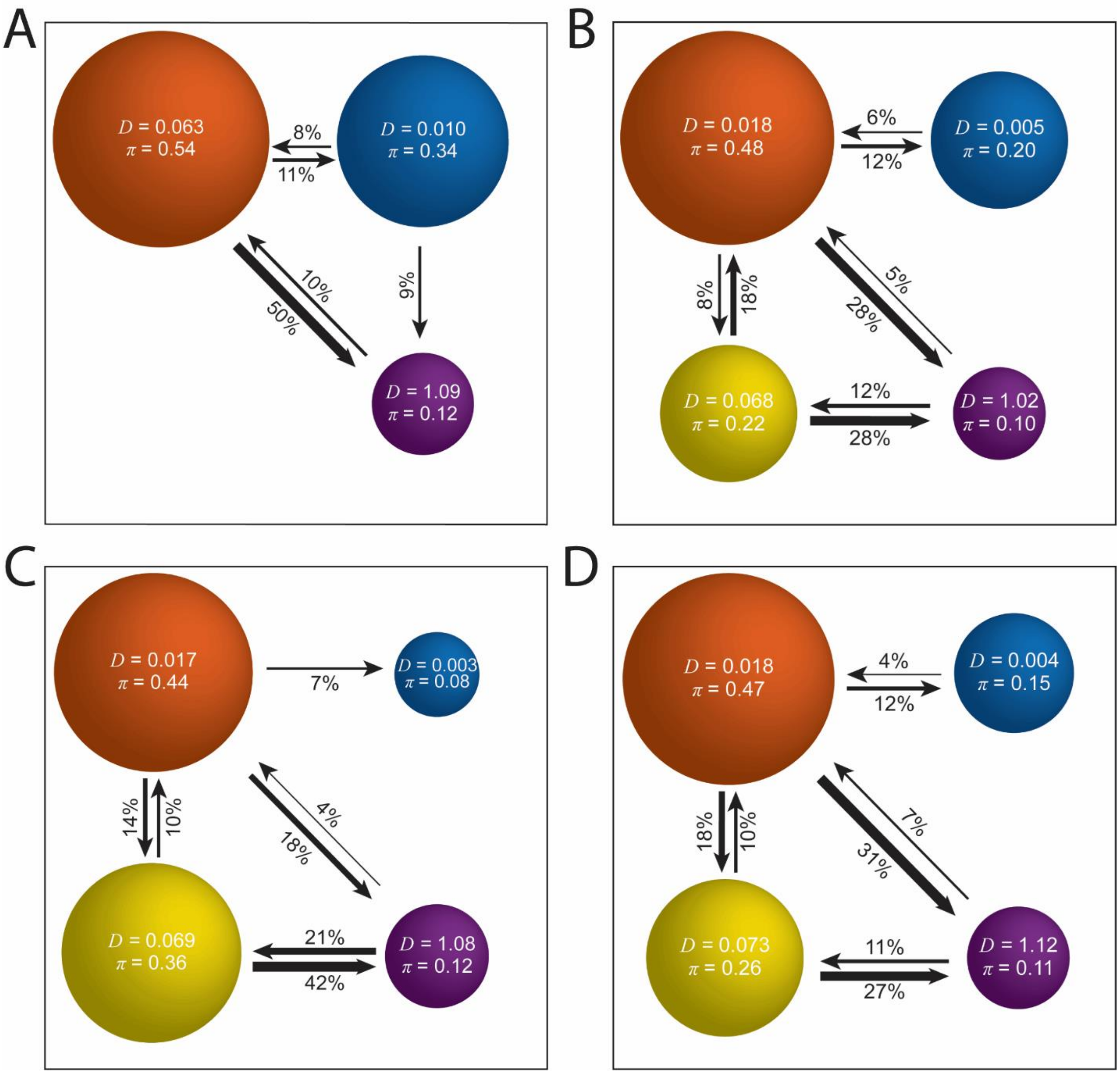


**Supplemental Figure 7:** TcpP-PAmCherry transition plots. Based on the identification of distinct diffusion states for TcpP-PAmCherry (circles with colors as in Figure 1C and with average single-molecule diffusion coefficient, *D*, indicated in $\mu m^2/s$), the average probabilities of transitioning between mobility states at each step are indicated as arrows between those two circles, and the circle areas are proportional to the weight fractions. Low significance transition probabilities less than 4% are not displayed. Numbers above the arrows indicate the probability of transition. A) *V. cholerae tcpP-PAmCherry toxTpro*∆(−55–+1), corresponding to main text Figure 2D. B) *V.*

*cholerae tcpP-PAmCherry* Δ*toxRS,* corresponding to main text Figure 3B*.* C) *V. cholerae tcpP-PAmCherry* Δ*toxRS toxTpro*Δ(−55–+1), corresponding to main text Figure 3D. D) *V. cholerae tcpP-K94E-PAmCherry,* corresponding to main text Figure 4B.

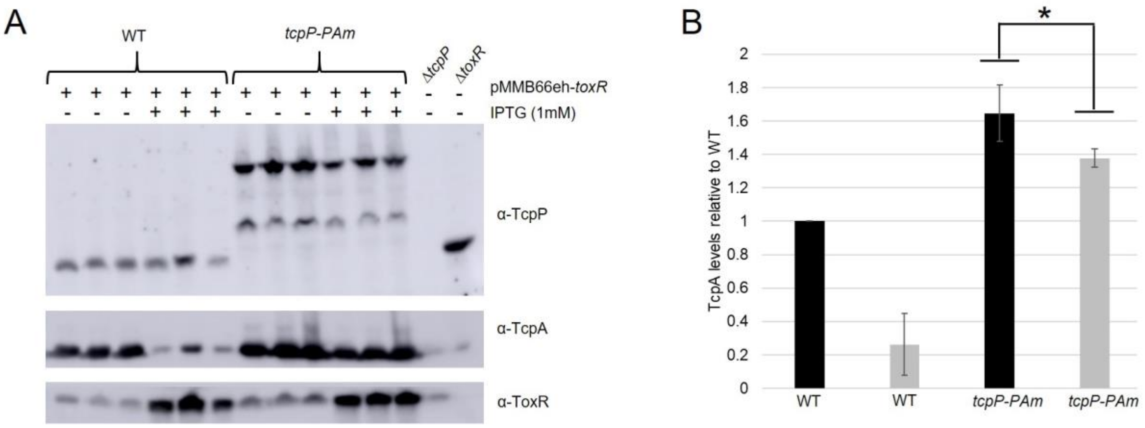


**Supplemental Figure 8:** ToxR overexpression reduces virulence factor production. A) Western blots of cell lysates, three biological replicates, collected after 6 hrs of virulence-inducing conditions with or without IPTG. B) Densitometry analysis of the TcpA western blot in panel A. ImageJ was used to perform the densitometry analysis. Black bars: −IPTG; grey bars: +IPTG. Error bars represent standard deviation. One-tailed Student's *t*-test was used to determine statistical significance. *indicates a P-value of 0.029.

**Supplemental Table S1:** Strain list.

| Strain | Description | Reference |
|---|---|---|
| *V. cholerae* 0395 classical biotype | Wild type | DiRita lab collection |
| *V. cholerae* Δ*tcpH* | Isogenic deletion | DiRita lab collection |
| *V. cholerae* Δ*tcpP* | Isogenic deletion | DiRita lab collection |
| *V. cholerae* Δ*toxRS* | Isogenic deletion | DiRita lab collection |
| *V. cholerae tcpP-PAmCherry* | Isogenic construct; TcpP-PAmCherry (C-terminal fusion), native *tcpH* start codon and 3rd amino acid mutated (ATG to GTG and AAA to TAA respectively), and both ribosomal binding site and coding sequence of *tcpH* cloned downstream of PAmCherry. | This study |
| *V. cholerae tcpP-PAmCherry* Δ*tcpH* | Isogenic construct | This study |
| *V. cholerae tcpP-PAmCherry* Δ*toxRS* | Isogenic construct | This study |
| *V. cholerae tcpP-PAmCherry* Δ*toxRS* toxTproΔ(−55–+1) | Isogenic construct | This study |

| | | |
|---|---|---|
| *V. cholerae tcpPK94E-PAmCherry* | Isogenic construct | This study |
| *V. cholerae tcpP-PAmCherry* toxTproΔ(−55–+1) | Isogenic construct | This study |
| *V. cholerae tcpP-PAmCherry* toxTproΔ(−112–+1) | Isogenic construct | This study |
| *E. coli* ET12567 Δ*dapA* | Cloning vector recipient | Allard, N., et. al. 2015. Canadian journal of microbiology, 61(8), pp.565-574. |
| *E. coli* ET12567 Δ*dapA* pKAS32-(empty vector) | Plasmid vector strain | DiRita lab collection |

**Supplementary Table S2:** Primer list. Kpn1-HiFi restriction sites were included in forward primers and Xba1 restriction sites were included in all reverse primers to provide homology between insert and vector sequences.

| Description | Sequence |
|---|---|
| pKAS-TcpP promoter FW | ctaacgttaacaaccggtacTTTCGAGTGATAGAAAAAGG |
| pKAS-TcpP FW | ctaacgttaacaaccggtacATGGGGTATGTCCGCGTG |
| TcpP-PAmCherry FW | atgcactaaaaatATGGTGAGCAAGGGCGAGGA |
| TcpP-PAmCherry RV | ccttgctcaccatATTTTTAGTGCATTCTAATGTCTTCTGTTC |
| TcpH-PAmCherry FW | ctaatgtcttCTTGTACAGCTCGTCCATGC |
| TcpH-PAmCherry RV | gctgtacaagAAGACATTAGAATGCACAAAAAATTAAAAG |
| Downstream TcpH-PAmCherry RV | tcatgataagaccCTTGTACAGCTCGTCCATGCC |
| Downstream TcpH-PAmCherry FW | cgagctgtacaagGGTCTTATCATGAGCCGCCTAG |
| pKAS-downstream TcpH RV | aaatttgcgcatgctagctatagttCTTGGTCTTTTTTAGATAACGTAAGC |
| TcpPK94E RV | GATCAACGTCTCATGTTCATC |
| TcpPK94E FW | GATGAACATGAGACGTTGATC |

| | |
|---|---|
| *toxTpro* Δ(−55–+1) RV | tcccaatcatATCTTAAAATCGAAGTTAATATAAAACTAC |
| *toxTpro* Δ(−55–+1) FW | gattttaagatATGATTGGGAAAAAATCTTTTC |
| pKAS-*toxTpro* Δ(−112–+1) FW | ctaacgttaacaaccggtacGTTGGTGGTGTTCCAGATAATAC |
| *toxTpro* Δ(−112–+1) RV | ttcccaatcaGTATTACATAAGAAAAACATAAAGTAACTCATG |
| *toxTpro* Δ(−112–+1) FW | tatgtaatacTGATTGGGAAAAAATCTTTTC |
| pKAS-*toxTpro* Δ(−112–+1) RV | tgcgcatgctagctatagttATCATCAGTAATAAATATAGAGTTATATTTTTTTTC |
| *recA* FW | ATTGAAGGCGAAATGGGCGATAG |
| *recA* RV | TACACATACAGTTGGATTGCTTG AGG |
| *toxT* FW | ACTGATGATCTTGATGCTATGGAG |
| *toxT* RV | CATCCGATTCGTTCTTAATTCACC |
